# Development of a Clinically Relevant Rabbit Model of Acute Laryngeal Injury

**DOI:** 10.1101/2025.06.27.662000

**Authors:** Ryan Stepp, Naushin Ali, Areli A. Rodriguez, Ethan Nicklow, Noah Thornton, Adithya Reddy, Hannah Kenny, Patrick S. Cottler, Donald R. Griffin, James J. Daniero

**Affiliations:** University of Virginia Department of Otolaryngology – Head and Neck Surgery, Charlottesville, VA 22908; University of Virginia Department of Biomedical Engineering, Charlottesville, VA 22908; Thomas Jefferson University Hospitals, Department of Otolaryngology-Head and Neck Surgery, Philadelphia, PA 19107

**Keywords:** Acute Laryngeal Injury, Laryngeal Trauma, Airway Stenosis, Fibrosis, Animal Model

## Abstract

**Objectives:** Acute Laryngeal Injury (ALgI) is created as a result of endotracheal tube pressure ulcer formation leading to fibrosis and inflammation. This condition often leads to airway obstruction and voice and swallowing dysfunction. This study demonstrates a reliable animal model of ALgI to reproduce the acute wound process seen clinically, to explore the pathophysiology of this disease process, and serve as a reproducible injury suitable for the evaluation of therapeutic interventions.

**Methods:** An ALgI model was developed in New Zealand White rabbits using precise mucosal stripping of the posterior larynx, followed by intubation with an oversized 4.0 endotracheal tube for one hour to mimic intubation-associated trauma and pressure ischemia. Laryngoscopy and laryngeal harvest were performed two weeks post-injury for histologic and immunofluorescent evaluation.

**Results:** Injured rabbits demonstrated an eight-fold increase in posterior glottic thickness (1.57mm vs. 0.19mm in controls; p=0.0004) and eleven-fold increase in collagen content (1.93mm^2^ vs. 0.17mm^2^; p=0.005). Collagen subtype analysis revealed a shift toward active collagen within the injured larynx compared to the uninjured, with increased type III collagen (69.0%% vs. 26.1%; p<0.0001) and reduced type I collagen (27.2% vs. 73.9%; p<0.0001) in the posterior glottis, consistent with the proliferative phase of wound healing. Collagen fiber alignment analysis demonstrated increased coherency in injured tissues (0.36 vs. 0.21; p=0.023), indicating early organized collagen formation consistent with scar formation within the posterior glottis.

**Conclusions:** The model offers a robust platform for studying the acute pathogenesis of laryngeal injury and for testing the treatment options in the management of ALgI.

**Level of Evidence:** Level III

## 1. Introduction

Acute laryngeal injury (ALgI) is a recognized complication of endotracheal intubation, especially in intensive care settings ^1–4^. Notably, recent clinical studies have shown that more than half of patients intubated for over 24 hours developed signs of ALgI, and a portion of these patients experience significantly worse breathing outcomes at three months post-extubation ^4^. This injury typically involves pressure trauma in the posterior glottis, specifically the cricoid, interarytenoid region and cricoarytenoid joints, where the endotracheal tube contacts the larynx ^1^. While often subtle at the time of extubation, ALgI can initiate a wound-healing cascade that progresses to mature fibrosis, leading to reduced glottic aperture and impaired vocal fold mobility ^1^ and results in dysphagia, hoarse voice, and vocal fold scar ^3^. In severe cases, long-term progression of ALgI fibrosis in the posterior larynx manifests as posterior glottic stenosis (PGS) – a condition characterized by scar-mediated narrowing of the glottic airway and fixation of the arytenoid cartilages, resulting in restricted vocal cord abduction and ventilatory compromise ^5^. PGS is a potentially life-threatening sequela of endotracheal tube trauma ^5^.

If the initial management of ALgI is unsuccessful and the fibrosis progresses to PGS, more invasive surgical interventions are often required to improve the airway. A multitude of surgical techniques have been described, including endoscopic laser excision of scar tissue, posterior cordotomy or partial arytenoidectomy, and open reconstruction with cartilage grafts or stents to prevent restenosis ^5^. Adjunct treatments such as steroid injection ^6^ or topical mitomycin-C ^7^ are sometimes employed to mitigate scar formation, but these do not consistently restore normal mucosa or synovial joint mobility. Indeed, PGS is well-known for requiring multiple procedures over time, as even successful initial management may be followed by scar contracture and re-stenosis that necessitates many repeat surgical treatments ^5^. Likewise, voice and swallowing outcomes after surgery can be variable since excising scar to relieve obstruction may further impair the delicate vocal mechanism.

Despite the frequency and morbidity of post-intubation laryngeal injuries, they have historically received little attention compared to other critical complications ^1^, and patients often present only after chronic symptoms (e.g. dyspnea or voice changes) arise 4-6 weeks after the initial injury ^8^. This underscores the need for greater awareness and research into the pathogenesis of ALgI and its progression to stenosis. Early intervention to treat ALgI to prevent the sequelae that leads to PGS have increased in importance to provide improved clinical outcomes ^1,7,9^.

A major obstacle to developing better treatments for acute laryngeal injury and fibrosis has been the lack of reliable animal models. Because the development of laryngeal stenosis involves complex, dynamic processes that cannot be ethically or feasibly monitored in humans, in vivo models are indispensable for controlled studies ^10^. However, to date there is no well-established, minimally invasive animal model that faithfully replicates the post-intubation acute laryngeal injury and its progression to fibrosis of the posterior glottis ^11^. This gap in experimental capability hampers the development of effective antifibrotic interventions for the larynx. Herein, we describe the establishment of a consistent animal model of fibrosis associated with ALgI and to histologically characterize its features, thereby providing a platform for investigating the pathogenesis of posterior laryngeal scar and evaluate treatment efficacy of the acute injury towards preventing PGS.

## 2. Methods

Ten New Zealand White rabbits were included for analysis, consisting of injured (n=5) and uninjured control (n=5) larynges. Animal experiments were performed under a protocol approved by the University of Virginia Institutional Animal Care and Use Committee (Animal Welfare Assurance A3245-01) in accordance with the National Institutes of Health’s Guide for the Care and Use of Laboratory Animals.

### 2.1. Acute laryngeal injury (ALgI) creation

At the initial injury time point (day 0), rabbits were anesthetized with an intramuscular injection of ketamine (50 mg/kg) and xylazine (5 mg/kg). The larynx was exposed using a custom rabbit-modified infant Parsons laryngoscope, placed in suspension with a ring stand and clamp. Photodocumentation was obtained using a 1.9 mm 0-degree Storz rigid telescope and Storz endoscopic camera and Storz AIDA digital capture device. Rabbits underwent mucosal stripping of the posterior glottis from one vocal process to the contralateral vocal process using cold biopsy forceps and microscissors, widely exposing the cartilage of each arytenoid and the intervening cricoid cartilage along the surface where an endotracheal tube would make contact. To mimic traumatic intubation, an oversized 4.0 endotracheal tube was inserted for one hour duration to further induce local pressure ischemia. Placement of an endotracheal tube of this size in a ∼3 kg rabbit provides resistance due to the snug fit and results in mildly traumatic intubation. During this time, rabbits were maintained on 2.0% inhaled isoflurane. After the procedure, rabbits were allowed to recover from anesthesia and were extubated without further intervention.

For pain management, each animal received pre-operative and 6-hour post-operative subcutaneous injections of buprenorphine (0.03 mg/kg). Additionally, a fentanyl dermal patch (25 μg) was applied for prolonged analgesia over 3 days.

### 2.2. Histology and immunofluorescence

At 14 days post-injury, laryngoscopy was performed to assess scarring and endoscopic images were collected. Rabbits were then humanely euthanized via intravenous overdose of pentobarbital. All 10 larynx specimens were harvested, immediately embedded in OCT compound, and flash-frozen in liquid nitrogen. Tissues were sectioned at 16μm using a cryostat (CryoStar NX50; ThermoFisher Scientific).

#### Histological staining

Hematoxylin and eosin (H&E), Picrosirius Red (PSR), Masson’s Trichrome, and immunofluorescence (IF) staining for collagen-specific analysis were performed (n = 5, ALgI and n = 5, uninjured) using previously established protocols for frozen tissues to evaluate histological changes and collagen composition in the posterior glottis. H&E and Masson’s Trichrome sections were imaged at 20x magnification. For PSR, imaging under polarized light was performed at 40x magnification. For IF, imaging was performed at 40x magnification. Primary antibodies included a mouse monoclonal antibody against collagen I (Sigma-Aldrich, Cat# C2456, Clone COL-1, 1:400 dilution in PBS) and a goat polyclonal antibody against collagen III (ProSci, Cat# 98-505, 1:200 dilution in PBS). Secondary antibodies were Donkey anti-mouse Alexa Fluor 647 (ThermoFisher, Cat# A-31571, 1:1000 dilution in PBS) for collagen I and Donkey anti-goat Alexa Fluor 488 (ThermoFisher, Cat# A-11055, 1:1000 dilution in PBS) for collagen III. Nuclei were counterstained using DAPI (1:1000 in PBS). All samples were imaged using a Leica DM6 upright microscope (Leica Microsystems GmbH, Germany).

### 2.3. Image Analysis

Histologic analysis of H&E, Masson’s trichrome, and collagen I and collagen III IF samples was performed using QuPath (version 0.5.1) ^12^ and analysis of PSR samples was performed using Fiji (ImageJ, Java 1.8.0_322) and the OrientationJ ^13^ plugin ^14^ image processing software.

#### Posterior glottic thickness

The thickness of the posterior glottis directly correlates with the level of stenosis and therefore, was considered the primary endpoint for the model. PGT was measured on H&E stained samples from the luminal surface to the edge of the posterior cricoid cartilage at three discrete, equidistant locations along the cricoid cartilage and averaged to obtain the mean thickness.

#### Collagen density

Masson’s trichrome stained samples were analyzed to determine the collagen density within the posterior glottis. The anterior portion was limited to two-thirds the length of the lumen from the anterior commissure. QuPath’s pixel classification was utilized to differentiate between collagen, muscle, nuclei, and background regions to quantify the total area occupied by collagen-positive pixels within the posterior glottis.

#### Collagen Type I and III quantification

Immunofluorescence staining was performed to quantify collagen types I, III within the posterior glottis. Quantitative analysis of collagen content was conducted in QuPath. The ROI for the posterior glottis was defined from the posterior cricoid and arytenoid cartilage to the level of lumen involvement. Pixel classification was generated, similar to the approach used for Trichrome analysis, to classify and quantify pixel regions as Type I collagen (red), Type III collagen (green), and co-localized Type I and III (yellow).

#### Collagen fiber alignment

Sections stained with PSR were imaged under polarized light to exploit the birefringent properties of collagen fibers. These sections were analyzed using Fiji (ImageJ) with the OrientationJ plugin to assess coherency of the collagen fiber orientation within the posterior glottis region similar to the IF analysis. Three discrete regions of along the posterior glottis (posterior commissure and bilaterally along the lateral lumen were analyzed. All images were acquired from a consistent field of view, capturing an approximate area of 4,000 µm^2^, to ensure comparability across samples. Fiber coherency (0 = random fiber alignment, 1 = complete fiber alignment) enables the assessment of organized versus disorganized scar architecture.

#### Statistical analysis

All statistical analysis was performed in GraphPad Prism 10. Statistical significance was fixed below a cutoff of 0.05. Values for various metrics were compared between injured and uninjured controls (normal). Measurements of posterior glottic thickness, collagen content, and fiber orientation were assessed using unpaired two-way t-tests. Collagen type was assessed using a two-way ANOVA and Bonferroni multiple comparison corrections.

## 3. Results

### 3.1. Acute laryngeal injury (ALgI)

ALgI’s were successfully created in the 5 experimental rabbits. Figure 1A shows an endoscopic image of an uninjured larynx, Figure 1B shows the creation of the injury with biopsy forceps and the exposure of the underlying cricoid cartilage, and Figure 1C shows the proliferative phase of wound healing with granulation tissue indicative of ALgI at 2 weeks post-injury.

**Figure 1:**
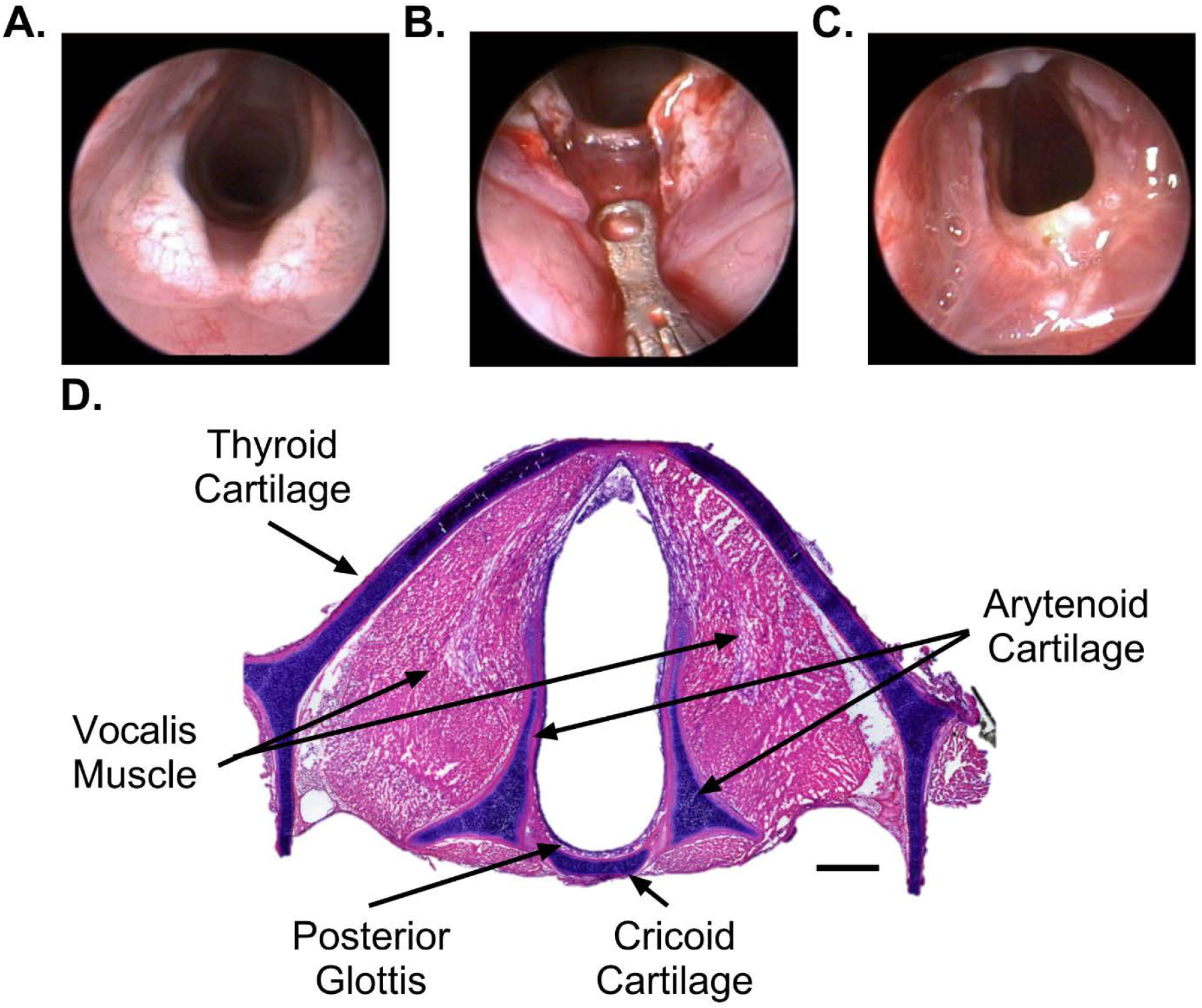
Endoscopic progression and anatomical orientation of the rabbit larynx and histologic landmarks for fibrosis quantification. (A) endoscopic visualization of the larynx pre-mucosal stripping, (B) mucosal stripping of the glottis to expose the cricoid cartilage, (C) two-weeks following injury with scar tissue development, (D) histological landmarks of larynx on H&E (scale bar = 1mm).

### 3.2. Histology and Immunofluorescence

#### Posterior glottic thickness

The evaluation of posterior glottis revealed a substantial hypertrophic response following injury (Figure 2A and B). At 14 days post injury, the PGT was significantly thicker in the ALgI group, 1.57 ± 0.55µm, compared to the uninjured normal group, 0.19 ± 0.05µm, (p = 0.0004) (Figure 2C).

**Figure 2:**
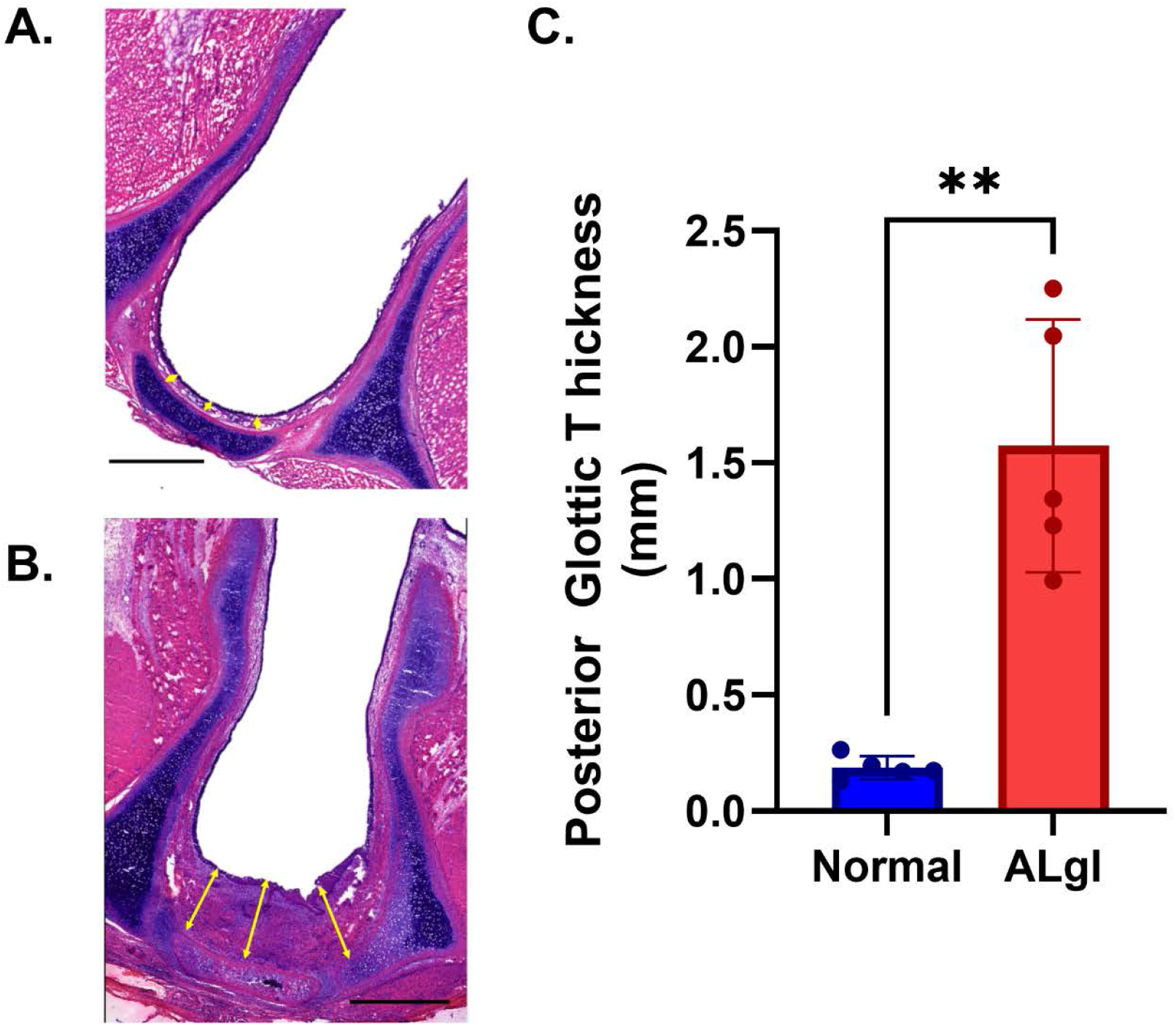
Posterior glottic thickness following acute laryngeal injury. (A and B) H&E-stained cross-section illustrating posterior glottic thickness in the uninjured and injured rabbit larynx respectively. Yellow overlaid lines marking the measured distance from luminal surface to posterior cartilage (scale bar = 1mm). (C) Average posterior glottic thickness measurements (** p<0.005).

#### Collagen density

Dense collagen content was seen in the scar compared to the normal glottis (Figure 3A and B). The total collagen area of nascent scar formation at the posterior glottis of the ALgI group, 1.93 ± 0.73mm^2^, was significantly greater than the uninjured group, 0.17 ± 0.08mm^2^, (p = 0.005). The difference reflects enhanced extracellular matrix deposition in the proliferative phase of healing prior to formal scar formation (Figure 3C).

**Figure 3:**
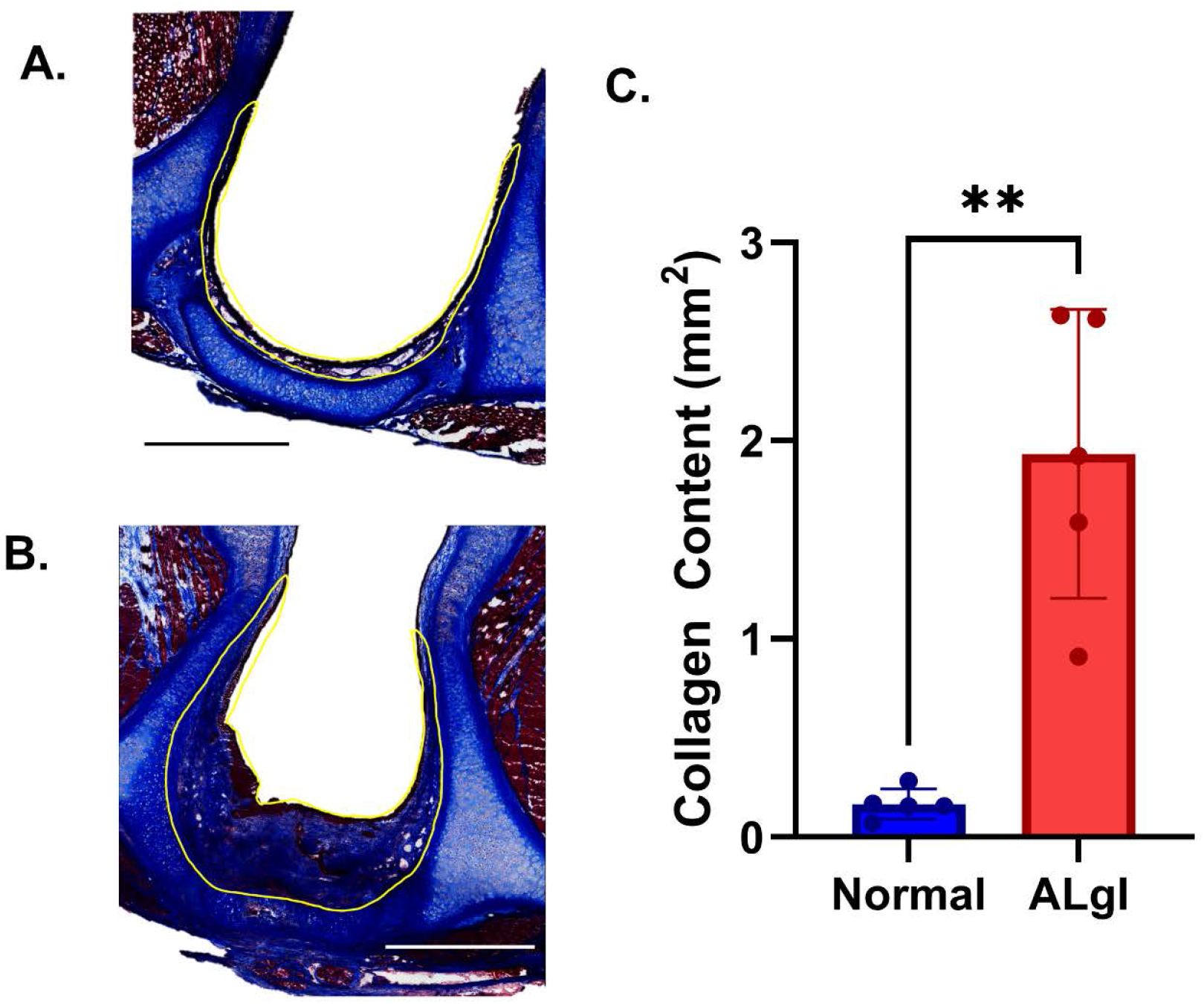
(A and B) Masson’s trichrome-stained cross-section illustrating posterior glottic thickness in the uninjured and injured rabbit larynx respectively. Yellow overlaid lines marking the glottic region of interest that was analyzed for collagen density (scale bar = 1mm). (C) Average area of collagen content within the region of interest as identified utlizing QuPath imaging software (** p<0.005).

#### Collagen Type 1 and III

Type I and Type III collagen was seen through IF within the posterior glottis in both groups (Figure 4A and B). The total Type I collagen within the scar of the ALgI group, 27.2% ± 5.1%, was significantly lower than within the uninjured glottis, 73.9% ± 13.3%, (p <0.0001) (Figure 4C). However, the total Type III collagen presence revealed the inverse, with 69.0% ± 3.3% in the injured group, compared to 26.1% ± 13.3% in the normal group (p < 0.0001) (Figure 4C). Areas of co-localized Type I and III collagen were significantly great in the uninjured group, 38.4% ± 19.5%, compared to 8.4% ± 7.1% in the ALgI group (p = 0.0014) (Figure 4C).

**Figure 4:**
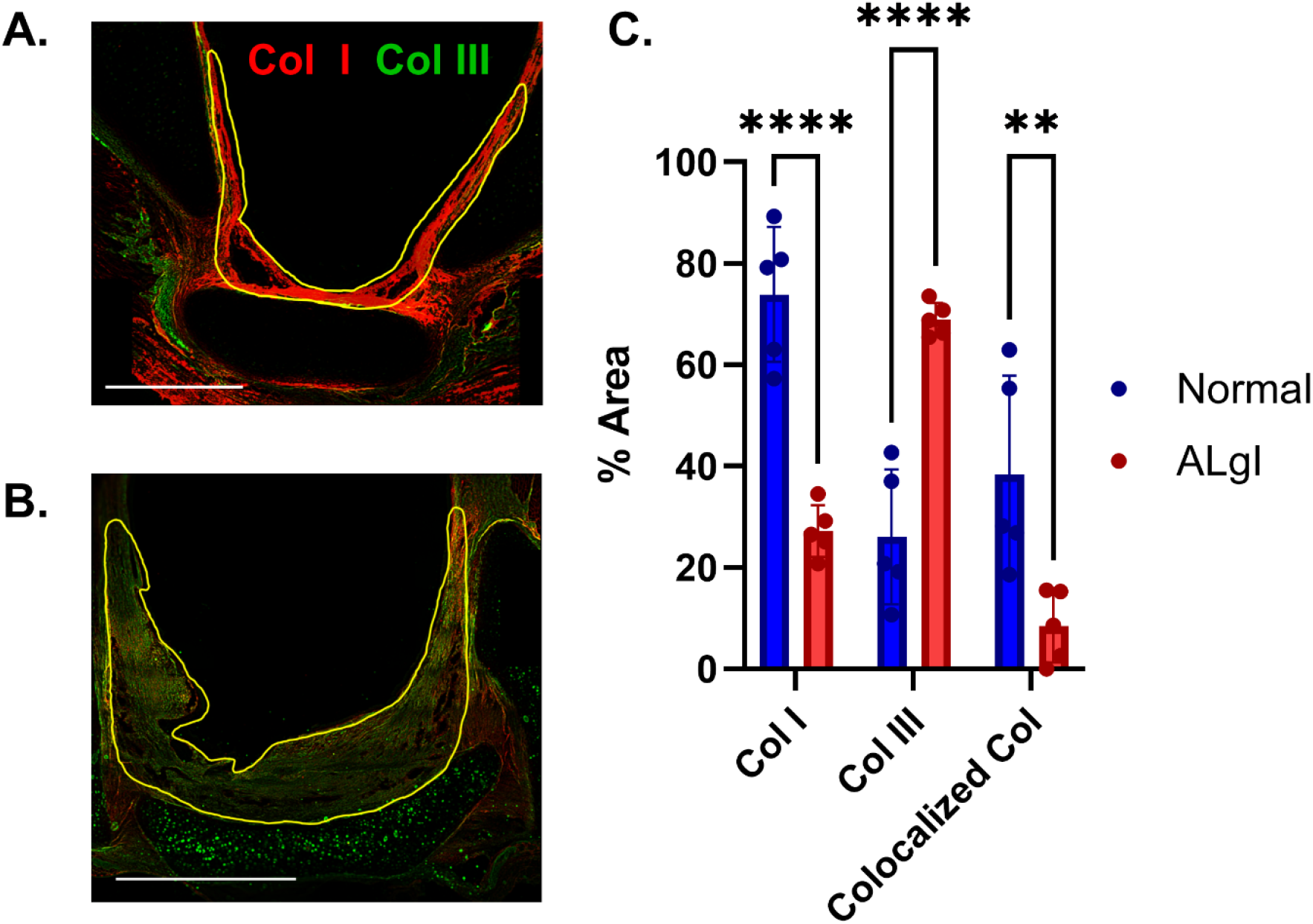
(A and B) Immunofluorescence of collagen type I and collagen type III within the posterior glottis in the uninjured and injured rabbit larynx respectively. Yellow overlaid lines marking the glottic region of interest that was analyzed for collagen type content (scale bar = 1mm). (C) Quantified levels of collagen type I, type III, and colocalized within the region of interest as identified utlizing QuPath image processing software (** p<0.005, **** p<0.0001).

#### Collagen fiber alignment

Fiber orientation was visualized through PSR staining in both the normal and injured posterior glottis (Figure 5A and B). Quantification of the alignment demonstrated significantly increased coherency in injured posterior glottic tissues, 0.36 ± 0.10, compared to the uninjured tissue, 0.21 ± 0.03 (p = 0.025) (Figure 5C). The increase in coherence indicates a more organized fiber architecture consistent with early scar tissue formation.

**Figure 5:**
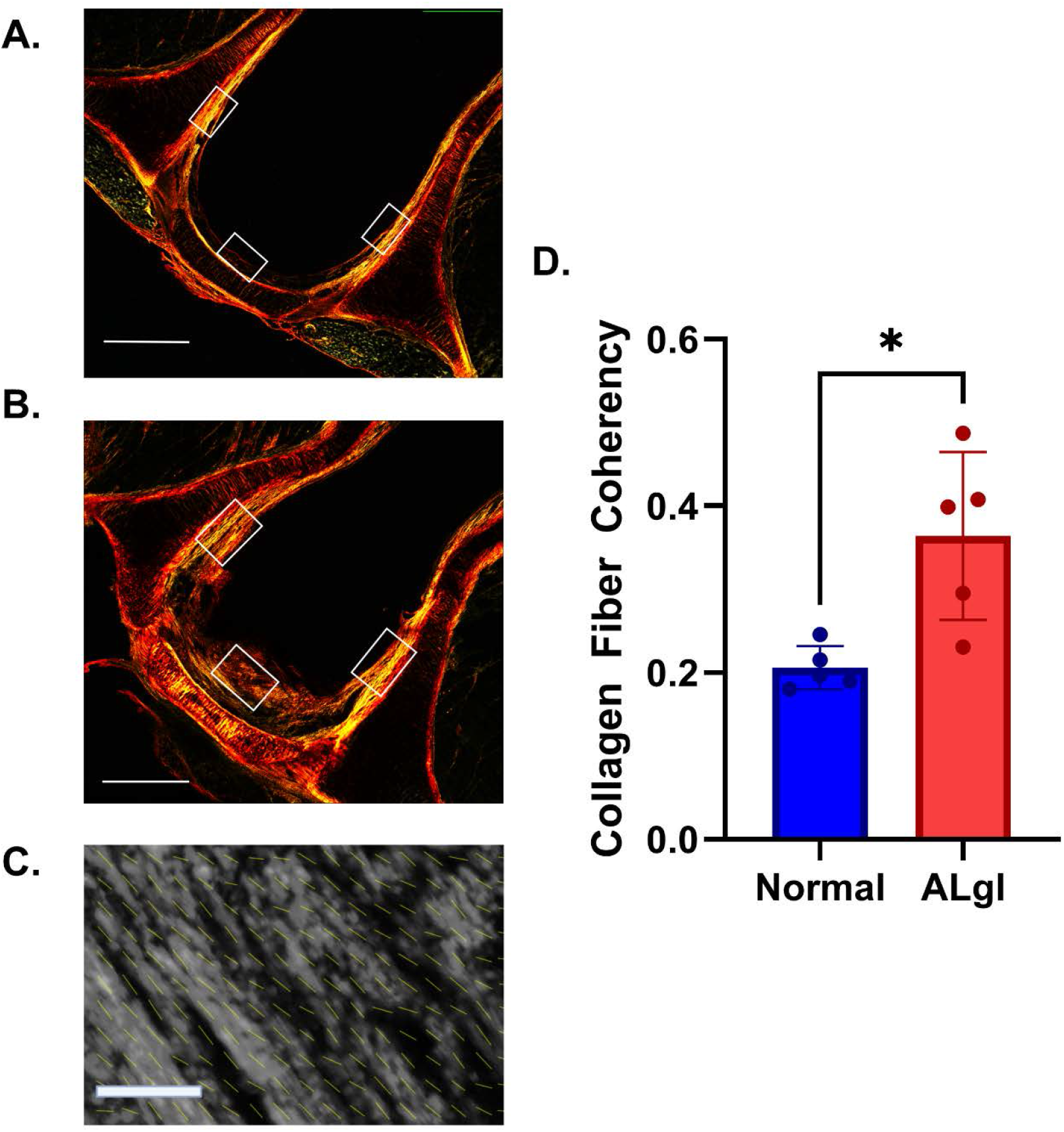
Collagen fiber alignment in posterior glottic scar tissue. (A and B) Picrosirus red stained images of the uninjured and injured rabbit larynx respectively imaged under polarized light. White boxes indicate the locations of 3 discrete regions of interest where collegen fiber orientation was calcualted (scale bar = 1mm). (C) Repesentative image of a region of interest with fiber coherency vector field map (scale bar = 20µm). (D) Quantification of the collagen fiber coherency caculated utilizing OrientationJ plugin for Figi image processing software (* p<0.05).

## 4. Discussion

The ALgI rabbit model presented here demonstrated hallmark features of laryngeal scarring, strongly mirroring the early clinical post-injury wound remodeling and early fibrosis. At 14 days post injury, endoscopic imaging clearly demonstrated the early scar formation associated with the acute injury. Quantitatively, injured larynges showed a significant increase in posterior glottic tissue thickness (mucosal and submucosal thickening), consistent with robust collagen deposition. Histologically, we observed a shift in extracellular matrix composition marked by elevated collagen content – particularly a predominance of collagen type III in the injured tissue. This finding differs from a previous model where vocal cord laser scarring was evaluated at 4 weeks ^11^, suggesting that in this model of ALgI, the two-week time point involves the development of early fibrosis and not mature scar ^15^. Moreover, collagen fibers in the injured posterior glottis became highly organized and aligned, evidence by increased fiber coherency measures. Such collagen fiber organization is a known hallmark of collagen evolving into more mature scar tissue in the larynx – scarred vocal folds, for example, develop densely packed, well-aligned collagen bundles compared to the random, random loose fiber network of normal mucosa ^16^. This structural remodeling underlies the stiffness of scarred laryngeal tissue and clearly recapitulated in our ALgI model. Taken together, the increases in glottic thickness, the collagen type III shift, and the higher collagen fiber alignment in our rabbits all validate that the model reliably produces a acute proliferative response and early fibrosis response that replicates what is observed in clinical laryngeal scar formation that often leads to PGS.

The consistency of our quantitative outcomes with those of prior laryngeal fibrosis models in animals further supports its translational relevance. Specifically, increased PGT following the acute injury is a consistent finding with the development of SGS ^17,18^. Tateya et al, 2006 also demonstrated early on Type III collagen development and later transition to Type I ^19^. Svistushkin et al, 2022, recognized collagen fiber organization as an important factor for “reduced scar index”, indicating restoration of a more normal collagen architecture in vocal fold regeneration ^20^. This was confirmed at 3 months, when effective treatments with mesenchymal stem cells led to collagen fiber to be more loosely packed and anisotropic ^21^. In sum, the ALgI model captures the transition from acute injury to early scar formation – a process that is characterized by mucosal ulceration, fibrin clot formation, and inflammation acutely. This is then followed by collagen-rich scar formation and wound contracture chronically – effectively mimicking the pathological progression seen in patients with laryngeal injury.

Though there are similar pathological features reported in other recent longer-term animal models of laryngotracheal stenosis, including exuberant fibroblast activity and thickened collagenous tissue in the injured larynx ^7^, this experimental model, focused on the acute injury, provides distinct differences and improvements to previous studies. A model of acute tracheal stenosis through prolonged endotracheal intubation (1 week) was achieved in a rabbit model by introducing the tube through a tracheostomy ^22^. However, the wound healing and subsequent scar formation associated with the invasive tracheotomy are complicating factors in survival and wound severity. Other large animal models focused on targeted induction of longer term PGS and not ALgI through laser ablation in canines ^23^ and rabbits ^11^, mucosal stripping ^24^, and brush abrasion ^25^ and did not include endotracheal intubation. Previous studies that utilized various protocols of intubation induced PGS led to non-standard levels of stenosis, making reproducibility difficult ^17,18,22,25,26^. Small animal models of an acute laryngeal injury have less clinical relevance due to the small tracheal size, and rely on mucosal damage without the inclusion of pressure induced ulceration ^19,27^.

Though we identified several histologic parameters that were statistically different between the injured and uninjured larynx, the small sample size (n=5/experimental group) is a limitation to the power of the study. In addition, the inclusion of the single 2 week time point, limits the detailed understanding of the progression of ALgI induced fibrosis.

A major motivation for developing a robust animal model of laryngeal injury is to expedite the discovery of treatments that can prevent or mitigate scar formation in the larynx and stop the progression to the more severe PGS. The presented ALgI rabbit model has already shown value in the testing of innovative therapeutic strategies. Our model opens the door for testing surgical interventions such as early endoscopic scar debridement, balloon dilation, or stent placement to prevent glottic closure, and non-surgical therapeutics, such as antibiotics, corticosteroids ^1,6^ biomaterials (e.g. injectable extracellular matrix gels ^28^ or slow-release anti-fibrotic carriers ^29^) aimed at restoring pliability and volume to scarred laryngeal tissues. In clinical practice, early steroid injection with or without balloon dilation after intubation injury has been attempted to reduce inflammation and fibrosis, but controlled data is limited due to life-threatening the nature of ALgI complications ^30^.

## 5. Conclusion

The analysis confirms significant differences in collagen deposition and posterior glottic thickness between normal and injured rabbits. These results validate the animal model for studying fibrosis and scar formation in ALgI, offering a basis for exploring therapeutic interventions aimed at mitigating fibrosis that can lead to PGS in such injuries. Future work into refining this model will include earlier time points as well as cellular and genetic profiles to provide mechanistic detail to the injury progression to guide treatment development.

## Acknowledgments

The authors would like to acknowledge Lisa Salopek, LVT, and Helen “Oli” Billcheck, LVT for animal care during this project

